# Acyl-CoA Thioesterases; a rheostat that controls activated fatty acids modulates dengue virus serotype 2 replication

**DOI:** 10.1101/2021.12.12.472314

**Authors:** Laura A. St Clair, Stephanie A. Mills, Elena Lian, Paul S. Soma, Aritra Nag, Caroline Montgomery, Gabriela Ramirez, Nunya Chotiwan, Rebekah C. Gullberg, Rushika Perera

**Author notes:** Address correspondence to Rushika Perera. College of Medicine, Medical University of South Carolina, Charleston, SC, USA. Department of Clinical Microbiology, Section of Virology, Umeå University, Umeå, Sweden (Nunya Chotiwan). Stanford University, Stanford, CA, 94305, USA. Laura A. St Clair and Stephanie A. Mills contributed equally to this work.

## Abstract

During infection with dengue viruses (DENVs), the lipid landscape within host cells is significantly altered to assemble membrane platforms that support viral replication and particle assembly. Fatty acyl-CoAs are key intermediates in the biosynthesis of complex lipids that form these membranes. They also function as key signaling lipids in the cell. Here, we carried out loss of function studies on acyl-CoA thioesterases (ACOTs), a family of enzymes that hydrolyze fatty acyl-CoAs to free fatty acids and coenzyme A, to understand their influence on the lifecycle of DENVs. Loss of function of the type I ACOTs 1 (cytoplasmic) and 2 (mitochondrial) together significantly increased DENV serotype 2 (DENV2) viral replication and infectious particle release. However, isolated knockdown of mitochondrial ACOT2 significantly decreased DENV2 protein translation, genome replication, and infectious virus release. Furthermore, loss of ACOT7 function, a mitochondrial type II ACOT, similarly suppressed DENV2. As ACOT1 and ACOT2 are splice variants, these data suggest that location (cytosol and mitochondria, respectively) rather than function of these proteins may account for the differences in DENV2 infection phenotype. Additionally, loss of mitochondrial ACOT2 and ACOT7 expression also altered the expression of several ACOTs located in multiple organelle compartments within the cell highlighting a complex relationship between ACOTs in the DENV2 virus lifecycle.

## Introduction

Dengue viruses (DENVs) are arthropod-borne viruses that are transmitted by the *Aedes aegypti* mosquito (1-2). These viruses infect over 400 million people each year. They are obligate intercellular parasites that rely on host cell metabolic pathways to fulfill their energy requirements and access substrates necessary for generating progeny virions (3). It is widely established that the lifecycle of DENVs is reliant upon host lipid metabolic pathways (3-6). Specifically, these viruses alter endoplasmic reticulum membranes to form scaffolds for viral protein translation and assembly of viral replication complexes (6,7). Moreover, during viral particle assembly, host cell membranes are co-opted and incorporated into the viral envelope as a structural component of the virus particle (3,6,7).

In our previous studies, we have shown that many lipid species are upregulated and are vital for the DENV serotype 2 (DENV2) lifecycle (8,9). Specifically, host phospholipids and sphingolipids are increased in abundance, some to benefit viral replication and others as a host response to infection (8-9, reviewed in 3). Precursors of these molecules are composed of fatty acyl-CoAs which are fatty acids that have been esterified to coenzyme A (CoA) (10,11). These fatty acyl-CoAs (activated fatty acids) can then undergo further modifications to be integrated into more complex lipids or be shuttled towards β-oxidation for cellular energy production (11).

Acyl-CoA thioesterases (ACOTs) are a family of hydrolases that control the intercellular balance between fatty acyl-CoAs and free fatty acids (FFAs). Specifically, they cleave fatty acyl-CoA into FFA and coenzyme A (10,11). There are 10 identified human ACOT enzymes further categorized into two groups by their domain motifs: α-β hydrolase (type I) and “hot dog” domain (type II) (Figure 1 A,B) (10,11). These enzymes are distributed throughout the organelles of human host cells including the cytoplasm, mitochondria and peroxisomes (Figure 1 A,B) (10). ACOTs have been implicated in the control of lipid metabolism by maintaining the ratios of fatty acyl-CoA and free fatty acids within each organelle (10,11). Upon perturbation, they have been shown to cause increased proliferation of cancer cells, are implicated in neurodegenerative diseases and play a protective role against diabetic cardiac damage (12-15).

**Figure 1.**
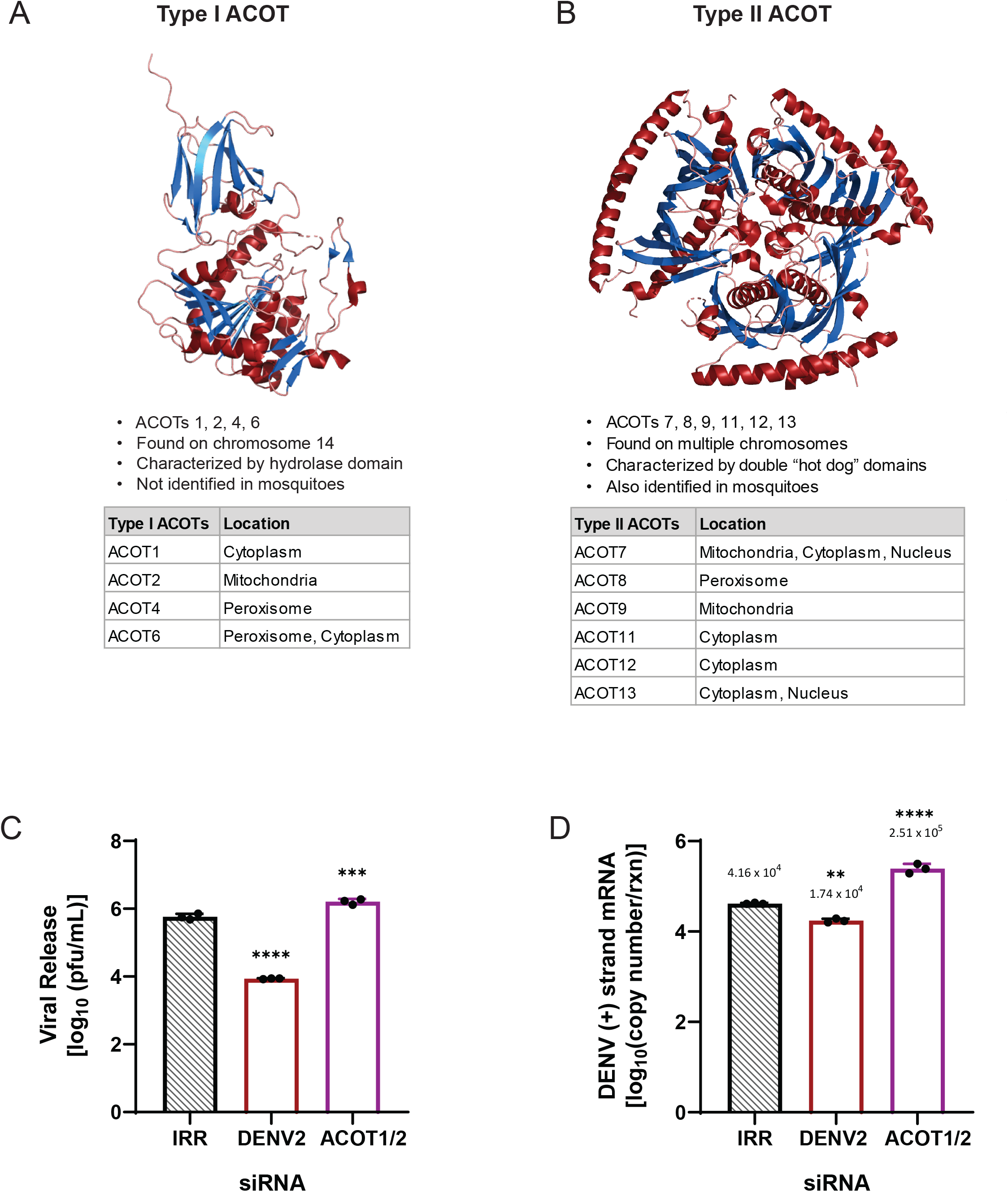
ACOT enzymes categorized by structural motifs and preliminary loss of function analysis of ACOT1/2. (A) Human ACOT2, Protein Data Bank identifier 3HLK is depicted. Type I ACOTs are characterized by a hydrolase domain that contains the active site. (B) Human ACOT7, Protein Data Bank identifier 2QQ2 depicted. Type II ACOTs contain the “hot dog” domain composed of two alpha helixes surrounding a hydrophobic core. Although the composition of the active site is unknown within the “hot dog” domain, these enzymes are functionally analogous to type I ACOTs (18). C-D: Huh7 cells were transfected with an siRNA pool targeting both the ACOT1 and ACOT2 genes as well as indicated controls and then infected with DENV2 for 24 hours (hr) (MOI = 3). (C) Infectious particle release was titrated via plaque assay. (D) Huh7 cells were collected, and relative copy number of viral RNA within cells was measured via qRT-PCR. qRT-PCR results were normalized to RPLP0. ACOT1/2: siRNA targeting acyl-CoA thioesterase 1 and 2, IRR: irrelevant siRNA control (no biological target for siRNA sequence), DENV2: siRNA targeting dengue virus, serotype 2. (A-B: images were generated utilizing PyMOL Molecular Graphics System, Version 1.2r3pre, Schrödinger, LLC. C-D: one-way ANOVA with Dunnett’s multiple comparisons test: * = p≤0.05, ** = p≤0.01, ***= p≤0.001, **** = p≤0.0001.)

Given that fatty acyl-CoAs are integral to lipid metabolism and energy homeostasis, we investigated if perturbing fatty acyl-CoA homeostasis by altering ACOT enzyme expression modulated DENV2 infection. We also investigated if the ACOT enzyme location (cytoplasmic vs. mitochondrial) and/or specificity type (type I vs. type II) differentially influenced DENV2 replication. We examined three representative ACOT enzymes, ACOT1, ACOT2, and ACOT7, to understand their impact on the lifecycle of DENV2 in human hepatoma (liver) cells (Huh7s). siRNA-mediated loss of function studies of the type I ACOTs 1 and 2 together significantly increased DENV2 infectious particle release. However, isolated knockdown of ACOT2 significantly decreased DENV2 protein translation, genome replication, and infectious virus release. Loss of function of ACOT7, a mitochondrial type II ACOT, also similarly suppressed DENV2. Furthermore, our studies reveal a complex relationship between type I and type II ACOTs during DENV2 infection that suggested a functional interdependency of these enzymes.

## Results

### Combined loss of ACOT1 and ACOT2 function increases DENV2 genome replication and infectious particle release

As ACOTs act as a rheostat for intracellular levels of FFAs and fatty acyl-CoAs, we hypothesized that ACOT functionality, which purportedly limits the availability of fatty acyl-CoAs, may have an inhibitory effect on the DENV2 lifecycle. To determine if ACOT enzymes affect DENV2 replication, we used siRNA-mediated knockdown to decrease the expression of type I ACOTs 1 and 2 in Huh7 cells. ACOT1 is located in the cytoplasm and ACOT2 in the mitochondria. There is ∼94% homology between the mRNAs of ACOT1 and ACOT2 with only an insertion of 90-110 nucleotides differentiating ACOT2 mRNA from ACOT1 mRNA (16,17,18). We included a non-target, irrelevant siRNA (IRR) to control for off-target effects of siRNA treatment and a DENV2-specific siRNA as a positive control for viral inhibition. We found that loss of ACOT1 and ACOT2 together resulted in an increase (∼178%) in infectious viral release (Figure 1C), and viral genome replication (∼470%, Figure 1D) compared to the IRR control. Due to the sequence similarity between ACOT1 and 2, we were unable to target siRNAs specific to ACOT1, but we were able to target specific siRNAs to ACOT2. Therefore, as shown below, we were able to parse out the influence of ACOT2 on this phenotype.

### Loss of ACOT2 function reduces DENV2 replication, infectious particle release and infectivity

Given the above observations from combined loss of ACOT1 and ACOT2 function, and that ACOT1 and ACOT2 share the same substrate specificity (18) we hypothesized that loss of ACOT2 function alone would also result in increased viral replication and release. Therefore, we repeated knockdown studies with siRNAs specifically targeting ACOT2 in Huh7 cells. Surprisingly, we found that loss of ACOT2 function resulted in a significant reduction (∼76%) in infectious virus release (Figure 2A) compared to the IRR control. Additionally, DENV2 genome replication was reduced by ∼50% following ACOT2 knockdown (Figure 2B). In our previous studies, we found that inhibition of specific enzymes involved in fatty acid metabolism resulted in the release of partially immature virions, thereby reducing particle infectivity (19). To determine if loss of ACOT2 function also reduced particle infectivity, we compared the ratios of viral RNA copies to infectious virions (particle/pfu ratio) in the supernatant of each of our treatment groups (Figure 2C). The data showed that there was a significant increase in the DENV particle/pfu ratio following ACOT2 knockdown compared to the IRR control; thus, suggesting a decrease in particle infectivity. We also found that viral protein translation was reduced by ∼50% (Figures 2D). We further confirmed that siRNA treatment was not cytotoxic to Huh7 cells (Figure 2E) and was also effective at reducing ACOT2 mRNA levels (Figure 2F). Overall, these data suggest that ACOT2 functionality is critical to the DENV2 lifecycle. This contrasts with the phenotype from the combined loss of function of ACOT1 and 2 suggesting that ACOT1 has a unique influence on viral replication.

**Fig 2.**
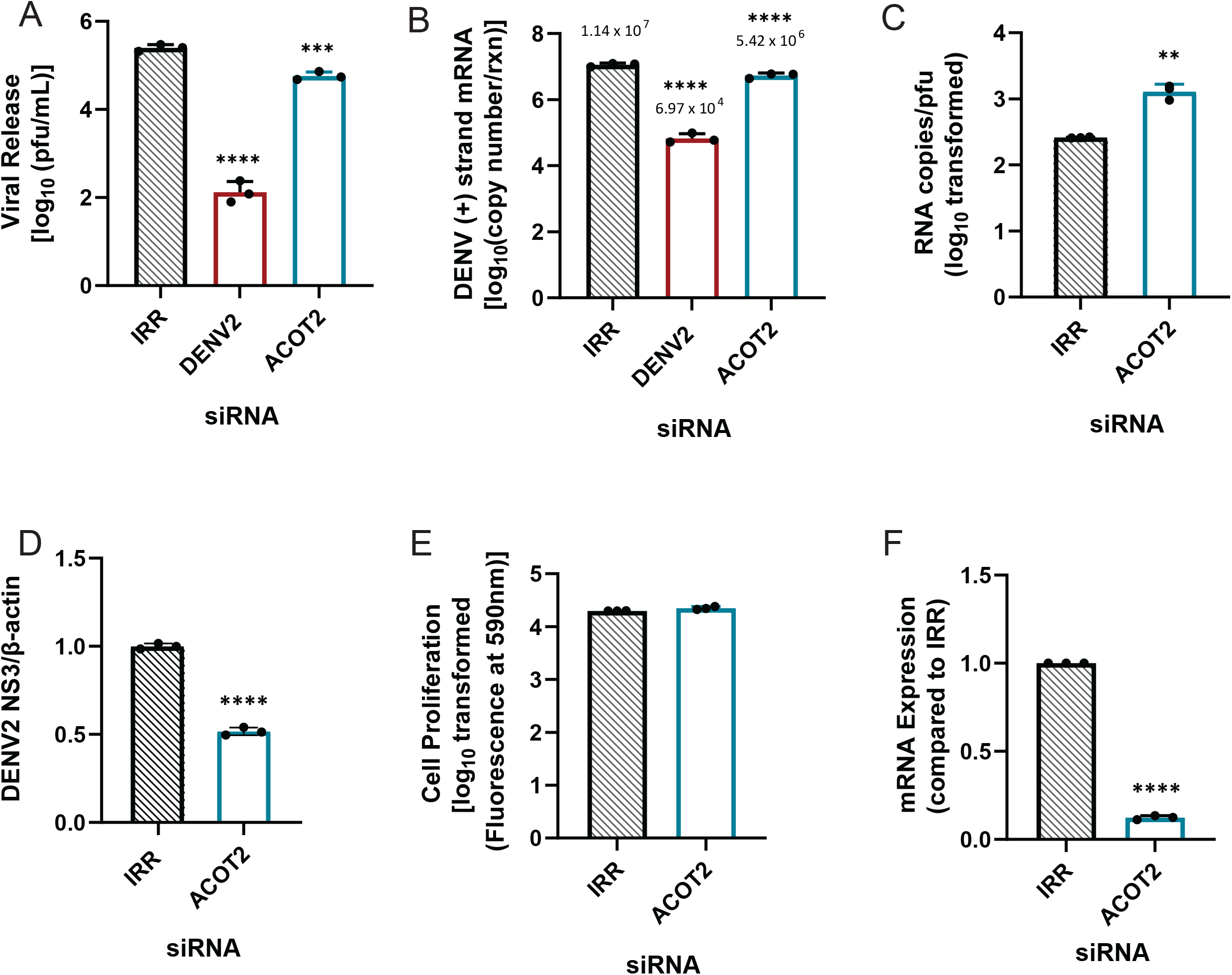
Loss of ACOT2 function reduces DENV2 genome replication and virus release. Huh7 cells were transfected with siRNA targeting the ACOT2 gene or indicated controls and then infected with DENV2 for 24hr (MOI = 0.3). (A) Infectious virus release was titrated via plaque assay. (B) Huh7 cells were collected, and relative copy number of viral RNA within cells was measured via qRT-PCR. Results were normalized to RPLP0. (C) Virus supernatant was collected at 24 hpi and split into two fractions. One fraction was titrated via plaque assay, while viral RNA from the other fraction was analyzed via qRT-PCR for total copy number of DENV2 positive-strand RNA. (D) Cell lysates were collected at 24 hpi, and analyzed via western blot. Samples were probed for DENV2 nonstructural protein 3, and normalized to β-actin. Li-cor IRDyes were used as secondary antibodies, and fluorescence intensity of each band was analyzed using area under the curve analysis in ImageJ. (E) A resazurin-reduction based cell viability assay was conducted to assess cytotoxicity of siRNA treatment. (F) Knockdown of ACOT2 mRNA was confirmed at 48 hr post transfection via qRT-PCR. ACOT2: siRNA targeting acyl-CoA thioesterase 2, IRR: irrelevant siRNA control (no biological target for siRNA sequence), DENV2: siRNA targeting dengue virus, serotype 2. (A-B: one-way ANOVA with Dunnett’s multiple comparisons test, C-F: unpaired t tests: * = p≤0.05, ** = p≤0.01, ***= p≤0.001, **** = p≤0.0001.)

### Mitochondrial ACOTs are vital for productive DENV2 infection

ACOT2 is suggested to be an important mediator of β-oxidation (20-21). This is the process of gaining ATP from the breakdown of fatty acyl-CoA to 2-carbon acetyl-CoA molecules. As this process occurs in the mitochondria, mitochondrial ACOTs may be important in regulating the balance of FFAs and fatty acyl-CoAs through shuttling fatty acyl-CoAs into β-oxidation (20-21). Previous studies indicate that β-oxidation is elevated during infection of host cells by DENVs (reviewed in 3). Both type I and type II ACOTs exist in the mitochondria. ACOT7 is a type II ACOT, and is functionally homologous to ACOT2 (both hydrolyze long-chain FAs) (18). Therefore, we investigated if loss of ACOT7 function also resulted in suppression of DENV2 infection. Similar to ACOT2 knockdown, we found that loss of ACOT7 function significantly reduced DENV2 infectious particle release (∼77%) in Huh7 cells (Figure 3A). Interestingly, ACOT7 inhibition resulted in a greater reduction of viral protein translation (∼75% reduction, Figures 3D) and viral genome replication (∼70% reduction, Figure 3B) compared to the effect of loss of function of ACOT2 (Figure 2B). Similar to the ACOT2 study, loss of ACOT7 function significantly decreased particle infectivity (Figure 3C). We confirmed that these results were also not due to cytotoxicity of the siRNA treatment (Figure 3E), and that the ACOT7 siRNA effectively reduced ACOT7 mRNA levels (Figure 3F). Taken together, these studies suggest that two of the mitochondrial ACOTs have a vital role in the DENV2 lifecycle.

**Fig 3.**
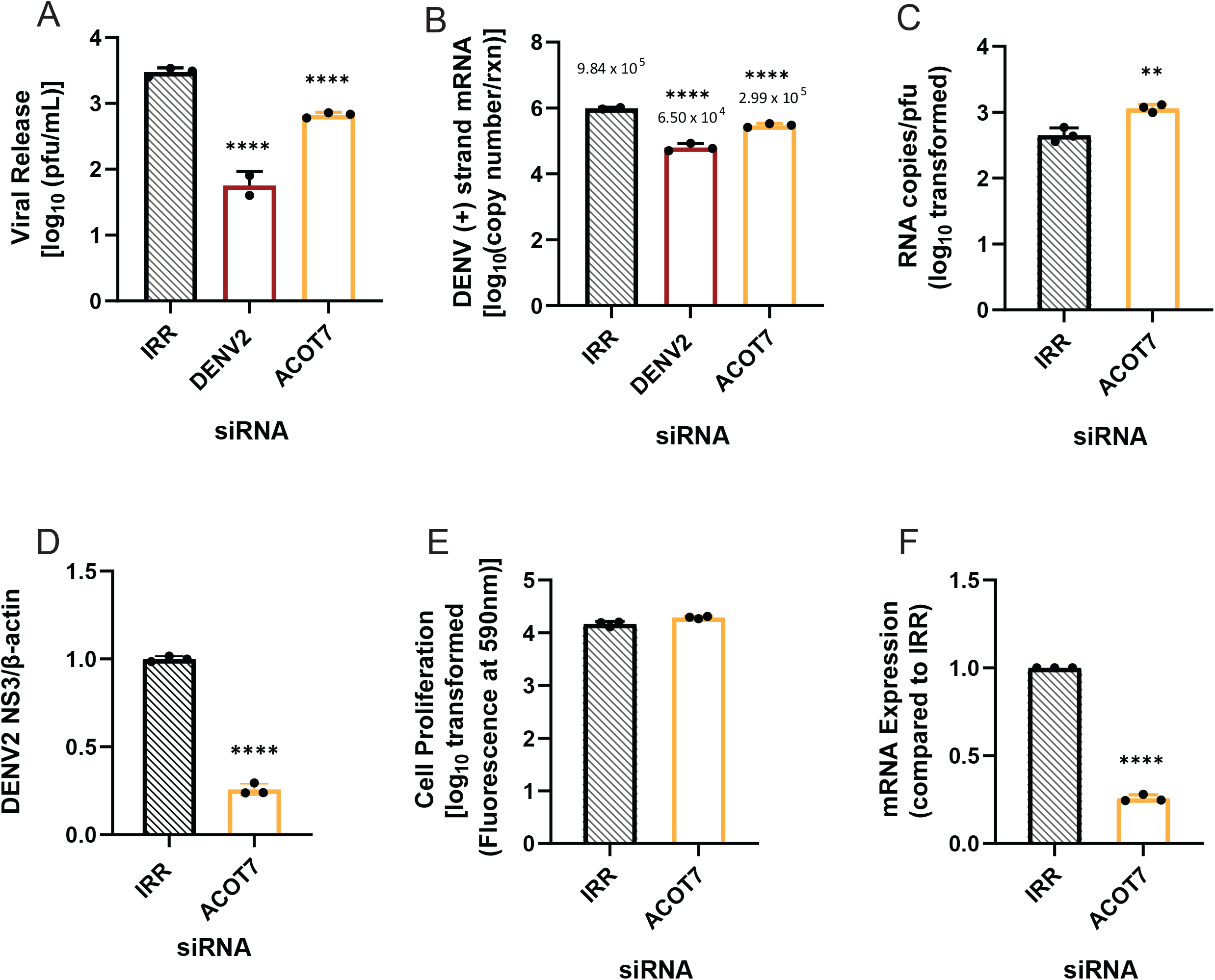
Loss of ACOT7 function suppresses DENV2 genome replication and infectious particle release. Huh7 cells were transfected with siRNA targeting the ACOT7 gene or the indicated controls and then subsequently infected with DENV2 for 24hr (MOI =0.3). (A) Infectious virus release was titrated via plaque assay. (B) Huh7 cells were collected, and relative copy number of viral RNA within cells was measured via qRT-PCR. Results were normalized to RPLP0. (C) Virus supernatant was collected at 24 hpi and split into two fractions. One fraction was titrated via plaque assay, while viral RNA from the other fraction was analyzed via qRT-PCR for total copy number of DENV2 positive-strand RNA. (D) Cell lysates were collected at 24 hpi, and analyzed via western blot. Samples were probed for DENV2 nonstructural protein 3, and normalized to β-actin. Li-cor IRDyes were used as secondary antibodies, and fluorescence intensity of each band was analyzed using area under the curve analysis in ImageJ. (E) A cell viability assay using resazurin was conducted to assess cytotoxicity of siRNA treatment. (F) Knockdown of ACOT7 mRNA was confirmed at 48 hr post transfection via qRT-PCR. ACOT7: siRNA targeting acyl-CoA thioesterase 7, IRR: irrelevant siRNA control (no biological target for siRNA sequence), DENV2: siRNA targeting dengue virus, serotype 2. (A-B: one-way ANOVA with Dunnett’s multiple comparisons test, C-F: unpaired t tests: * = p≤0.05, ** = p≤0.01, ***= p≤0.001, **** = p≤0.0001.)

### Both Type I and Type II ACOTs are differentially expressed upon ACOT2 or ACOT7 knockdown

Currently, there are ten human ACOTs known to exist in different subcellular compartments, and they have substrate specificity for a wide range of fatty acyl-CoAs (17,18). However, the functional overlap and/or ability of each ACOT to compensate for loss of function of the others is unknown. As mitochondrial ACOTs seem to play vital roles in fatty acid metabolism and energy production, we determined whether other ACOTs could compensate for loss of ACOT2 or ACOT7 function, both within and without the context of DENV2 infection. To investigate this, we carried out similar siRNA-mediated knockdown of ACOT2 and ACOT7 in both mock-infected and DENV2-infected Huh7 cells. An IRR siRNA control was also included. 24 hours post infection (hpi), mock- and DENV2-infected cells were collected and cellular mRNA levels of each ACOT enzyme was determined via qRT-PCR.

In both mock-infected and DENV2-infected samples, we observed similar trends in the mRNA expression of other ACOTs upon knockdown of ACOT2 (Figure 4 A,B) or ACOT7 (Figure 4 C,D). Specifically, we observed that in both mock- and DENV2-infected Huh7 cells, knockdown of ACOT2 significantly reduced mRNA expression of ACOTs 1, 6, 8, and 11 (Figure 4 A,B). A decrease in ACOT9 and ACOT12 mRNA expression was only observed in DENV2-infected, ACOT2 knockdown samples. We found loss of ACOT7 significantly reduced mRNA expression of ACOTs 2, 4, 8, 9, and 11 in both mock- and DENV2-infected cells, (Figure 4 C,D). Additionally, in DENV2 infected cells, ACOT6 and ACOT13 mRNA expression was decreased in ACOT7 siRNA-treated cells. It should be noted that the siRNAs for ACOT2 and ACOT7 do not have any sequence similarity to the other ACOT mRNAs. Therefore, these data suggests that other ACOTs may be functionally dependent on the activity of ACOT2 and 7. Intriguingly, inhibiting ACOT7 also reduced expression of ACOT2 (Figure 4 C,D); however, inhibition of ACOT2 did not impact the expression of ACOT7 (Figure 4 A,B). Thus, while both mitochondrial ACOTs are functionally similar, neither compensates for loss of the other at a transcriptional level, and ACOT2 functionality may rely upon ACOT7.

**Fig 4.**
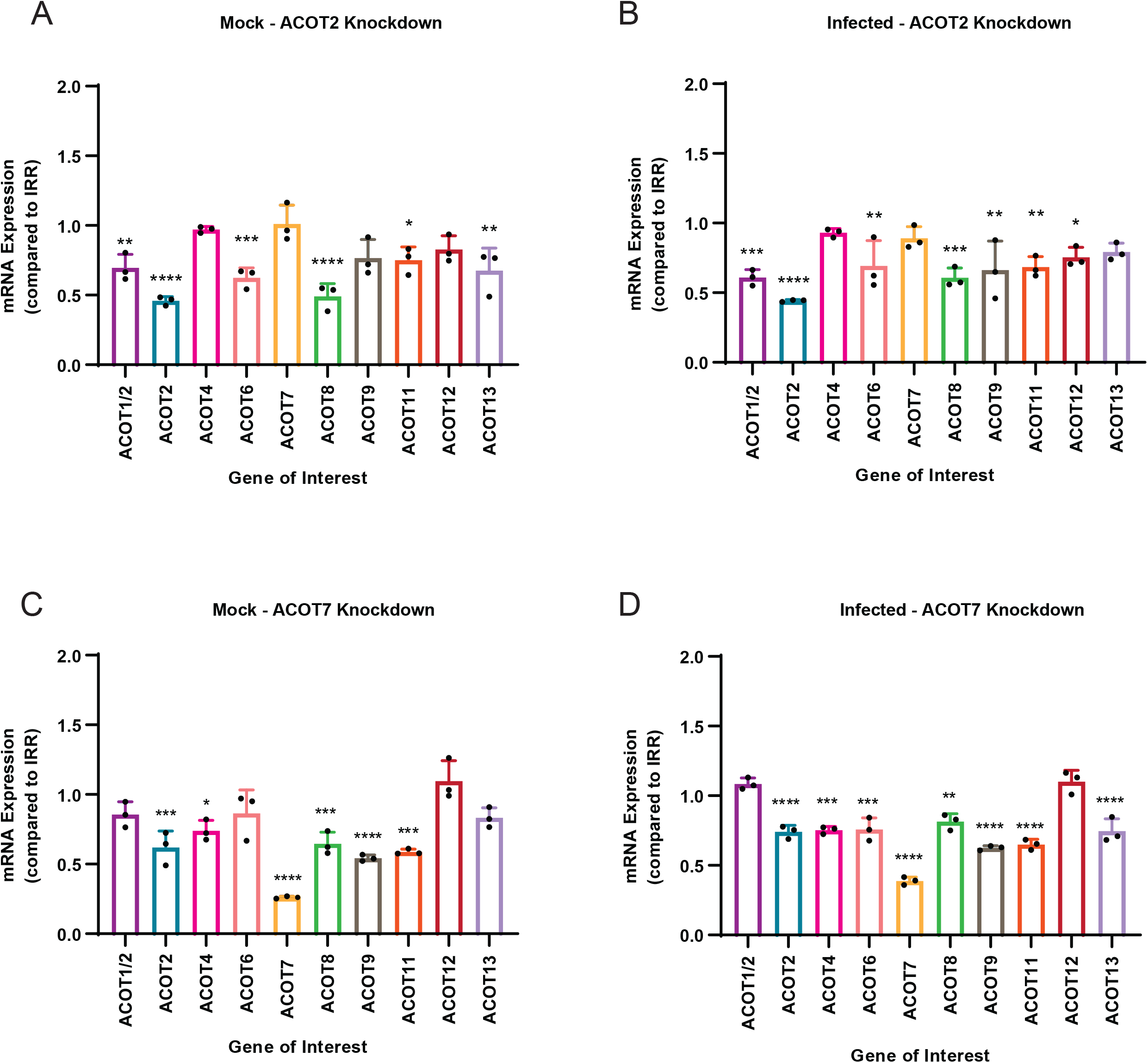
Inhibition of mitochondrial ACOTs underlines importance of mitochondrial ACOT functionality for expression of other ACOTs. Huh7 cells were treated with either ACOT2 (A,B) or ACOT7 (C,D) siRNA or an IRR control siRNA, and then (A,C) mock-infected or (B,D) DENV2-infected (MOI = 0.3). At 24 hpi, cells were collected and mRNA levels of all 10 human ACOTs was determined via qRT-PCR. (A-D: one-way ANOVA with Dunnett’s multiple comparisons tests, * = p≤0.05, ** = p≤0.01, ***= p≤0.001, **** = p≤0.0001.)

### Mitochondrial ACOTs are upregulated at early timepoints of infection

In previous studies, it has been demonstrated that DENV2 infection results in both viral- and host-mediated modulation of enzymes involved in fatty acid metabolism (3-9, 19). As our loss of function studies of ACOT2 and ACOT7 indicated these enzymes are vital for the DENV2 lifecycle, we analyzed whether ACOT2 and ACOT7 mRNA expression was altered over a time course of DENV2 infection. For this study, we collected both mock-infected and DENV2-infected cells at 0, 6, 24, and 48 hpi. These time points represent early, peak, and late viral replication. ACOT mRNA expression was analyzed using qRT-PCR (Figure 5). We observed that ACOT2 and ACOT7 mRNA expression was significantly upregulated at 6 hpi in DENV2-infected cells, but downregulated at 24 and 48 hpi (Figure 5A). A similar trend was noted when we compared mRNA expression between DENV2 and mock-infected samples at each timepoint although the increased expression at 6 hpi was not statistically significant (Figure 5B). These results combined with our results in Figures 2-3, suggest that ACOT2 and ACOT7 functionality is required for early stages of the DENV2 life cycle.

**Fig 5.**
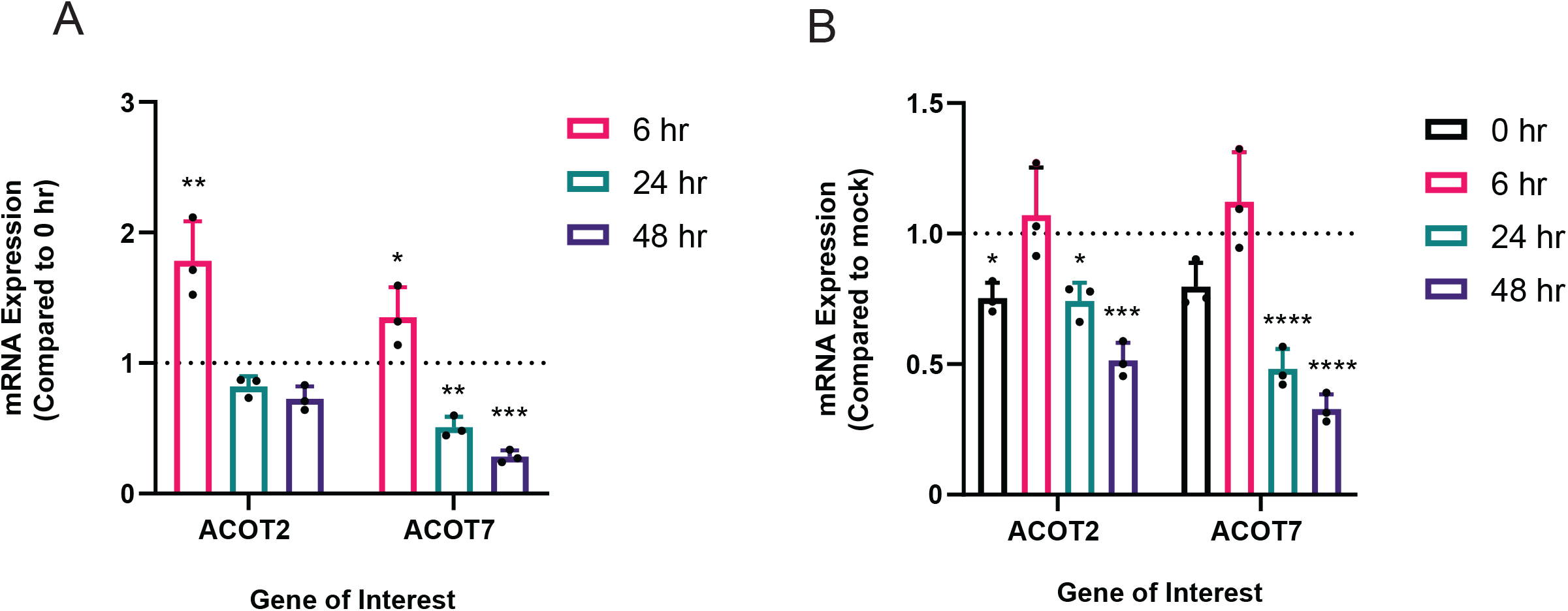
Mitochondrial ACOT mRNA expression is temporally altered during DENV2 infection. Huh7 cells were either mock-infected or DENV2-infected (MOI = 10). At 0, 6, 24, and 48 hpi, cells were collected and mRNA levels of ACOT2 and ACOT7 were assessed via qRT-PCR. DENV2-infected samples collected at 6, 24, and 48 hpi were compared either to (A) the 0 hpi DENV2-infected samples or to (B) mock infected samples at matched timepoints. Results are reported as an expression ratio (A) between each timepoint sample and the 0 hpi sample in DENV2-infected cells, or (B)between each timepoint sample and its respective mock-infected timepoint. (A-B: one-way ANOVA with Dunnett’s multiple comparisons test: * = p≤0.05, ** = p≤0.01, ***= p≤0.001, **** = p≤0.0001.)

## Discussion

The ACOT family of enzymes are suggested to be key regulators of the intercellular balance of activated fatty acids (fatty acyl-CoAs) and FFAs (10, 18). As previous studies have shown that DENVs are reliant upon activated fatty acids for completion of their lifecycle (8-9, 19, 22-23, reviewed in 3). Our present study investigated whether ACOT enzymes may be important modulators of DENV2 infection. The ACOT enzymes chosen represented both type I (ACOT1 and ACOT2) and type II ACOTs (ACOT7) as well as ACOTs found within the same organelle (ACOT2 and ACOT7 in the mitochondria). Interestingly, we found that loss of function of these enzymes had a divergent effect on the DENV2 lifecycle that was independent of their substrate specificity, and, instead, highlighted a more nuanced relationship between DENV2 and the subcellular regulation of fatty acid metabolism. Specifically, we found that the loss of function of the cytoplasmic ACOT1 enzyme resulted in increased DENV2 genome replication and infectious virus release, indicating that it’s normal function may be antiviral. However, loss of function of the mitochondrial ACOT2 and ACOT7 enzymes resulted in suppression of multiple stages of the DENV2 lifecycle indicating that the function of these enzymes is vital for effective DENV2 infection. Furthermore, our analyses revealed a functional dependency between enzymes within the ACOT family.

Because ACOTs act directly on activated fatty acids, it is predicted that they play a significant role in modulation of fatty acid metabolism (10, 11, 16, 18, 25). However, mechanistic insight as to the specific functions of ACOT enzymes is limited. One challenge is that the ester bond that links coenzyme A with fatty acids is labile which limits the ability to obtain a true ratio of activated fatty acids and FFAs in cells with traditional methods (10, 21). Thus, much of what the field understands about the function of these enzymes is concluded from studies characterizing their biophysical and biochemical properties, or from mouse models studying the impact of ACOT gene deletion on cardiac, neurological, and metabolic disorders (10-18, 20, 21, 26). During curation of this paper, we successfully developed an LC-MS assay that allowed us to distinguish between arachidonic acid, SH-CoA, and arachidonoyl-CoA (the primary substrate of ACOT7, discussed in 18 and 21). However, due to limitations in scalability of siRNA treatment, levels of these metabolites were below our limits of detection. As a future direction, use of knockdown cell lines may further characterize ACOT enzyme functionality. Moreover, an additional challenge was that ACOT1 and ACOT2 enzymes share 98% similarity at the protein level, and 94% nucleotide sequence homology (11, 17). This inhibited differentiation of these enzymes at the protein level. However, we were able to design specific siRNA and primer sequences that targeted the mitochondrial localization sequence of ACOT2, allowing us to characterize these enzymes based on their mRNA expression.

In an *ACOT1* knockout mouse model, ACOT1 was shown to modulate liver fatty acid metabolism during fasting, and loss of function of this enzyme led to increased triglyceride turnover and beta oxidation (16). Importantly, infection with DENV2 mimics a fasting state in the cell, and previous studies have established that triglyceride hydrolysis and beta-oxidation are increased in DENV2-infected cells (22, 24). Thus, inhibition of ACOT1 may further enhance these processes, leading to a more favorable environment for DENV2 replication. Mitochondrial functional assays have suggested that ACOT2 and ACOT7 functions to decrease beta-oxidation overload, possibly by mediating efflux of fatty acids from the mitochondrial matrix (21, 25, 26). Thus, loss of ACOT2 and ACOT7 function may have resulted in a buildup of fatty acyl-CoAs levels in the mitochondria, decreased turnover of CoA and fatty acid precursor molecules, and increased oxidative stress – all of which would antagonize the DENV2 lifecycle. Interestingly, we found that following ACOT2 or ACOT7 knockdown there was no compensation by any of the other mitochondrial ACOTs (ACOT2, 7 or 9). However, ACOT2 and ACOT7 knockdown decreased peroxisomal ACOT4, ACOT6 and ACOT8 expression. Peroxisomal ACOTs are important in degradation of very long-chain fatty acids that cannot be directly shuttled to the mitochondria (10,11). Peroxisomes are becoming recognized as important mediators for controlling or facilitating virus infection, including antiviral immune responses, interactions with viral capsid proteins, and influencing membrane fluidity (30-31). Therefore, another possibility is that ACOT2 or ACOT7 knockdown is indirectly suppressing DENV2 replication through altering expression of peroxisomal ACOTs which may impact peroxisomal lipid degradation activity. The role of peroxisomes in these biochemical interactions between virus and host is yet an uncharted territory.

In summary, we found there is a differential impact of ACOTs on DENV2 genome replication and infectious particle release that could be influenced by the subcellular location of these enzymes. We also observed that loss of function of a single ACOT impacted the expression of multiple ACOTs in different cellular locations highlighting the complexity of understanding how ACOTs influence viral replication. Overall, our current study underscores that DENV2 is reliant upon careful coordination of fatty acid metabolism to complete its lifecycle. Future studies aimed at characterizing the exact substrate specificities would increase our understanding of how the loss of function of specific enzymes regulate infection. Additionally, our data suggest that determining the functional relationship between ACOTs is warranted.

## Materials and Methods

### Cell lines and viruses

The cell lines used in this study were as follows: Human hepatoma cells (Huh7) (unknown sex, From Dr. Charles Rice) (32), Clone 15 (ATCC CCL-10) of the Baby Hamster Kidney Clone 21 cells (BHK-21), and C6/36 cells (ATCC CRL-1660, larva, unknown sex). Huh7 cells were maintained in Dulbecco’s Modified Eagle Medium (DMEM) (Gibco, LifeTech) while BHK-21 and C6/36 cells were maintained in Minimum Essential (MEM) (Gibco, LifeTech). All culture media was supplemented with 2 mM L-glutamine (HyClone), 2 mM nonessential amino acids (HyClone), and 10% fetal bovine serum (FBS) (Atlas Biologicals). C6/36 media was also supplemented with 25 mM HEPES buffer. Cells were incubated at 37°C with 5% CO_2_.

For this study, dengue virus serotype 2 was used (DENV2, strain 16681) (33-34). The virus was passaged in C6/36 cells. Viral titer was determined via plaque assay on BHK-21 cells as previously described (35). Virus infections were carried out at room temperature for 1 hour (hr), allowing for viral adherence. Subsequently, virus was removed, and cells were washed with 1xPBS before addition of media supplemented with 2 mM nonessential amino acids, 2 mM L-glutamine, and 2% FBS. Cells were incubated at 37°C with 5% CO_2_ for 24 hours.

### siRNA transfection and knockdown confirmation

ACOT1/2, ACOT2 and ACO7 loss of function was conducted by transfecting Huh7 cells with pooled siRNAs (Horizon Discovery/Dharmacon, Sigma-Aldrich – see Supplementary Table 1) as described previously (23) using RNAiMAX (Invitrogen) and incubating for 48 hr at 37°C with 5% CO_2_. At 48 hr post transfection, cells were either collected for a cytotoxicity assay (described below), knockdown confirmation or infected with DENV2 (as described above). At 24 hpi, cells and viral supernatant were collected for further analysis. Viral titration was completed via plaque assay. RNA was extracted from cells, and qRT-PCR analysis was used to confirm knockdown of ACOT2 and ACOT7 mRNA transcripts. Cellular transcripts were normalized to RPLP0, and the ACOT2 and ACOT7 siRNA-treated samples were then compared to an irrelevant control (IRR) using the delta delta cq method described in (19). Cytotoxicity of siRNA treatment was determined using a 1:10 dilution of resazurin (ThermoFisher) in cell culture media, and incubating cells for 1-2hr. Fluorescence was read on a Victor 1420 Multilabel plate reader (Perkin Elmer) at 560nm/590nm (excitation/emission).

### RNA extraction and qRT-PCR

RNA was extracted from cells and viral supernatant using TRIzol or TRIzol LS (ThermoFisher), respectively, following standard TRIzol extraction methods. For qRT-PCR, the Brilliant III Ultra-Fast SYBR® Green one-step qRT-PCR kit (Agilent) was used, and all reactions were set up according to manufacturer’s protocols on a LightCycler 96-well real-time PCR machine (Roche). The following cycling parameters were employed: 20 mins at 50°C for reverse transcription, 5 mins at 95°C, followed by 45 two-step cycles of 95°C for 5 secs, and 60°C for 60 secs. A melt curved followed each step starting at 65°C and ending at 97°C. (Primer sequences are reported in Supplementary Table 1). A standard curve was generated using *in vitro* transcribed viral RNA from a DENV2 cDNA subclone (derived from strain 16681 full-length clone) in order to quantify DENV2 genome copies. All cellular RNA transcript copy numbers were normalized to Ribosomal Protein Lateral Stalk Subunit P0 (RPLP0) RNA using the delta delta cq method as described previously (19, 36).

### Western Blotting

Huh7 cells were lysed in RIPA Buffer. Total protein in each sample was measured using the Qubit BR Protein Assay (ThermoFisher) on a Qubit Flex Fluorometer. Equal total protein was loaded from each of the indicated cell lysates and run on a Criterion^™^ XT 4-12% Bis-Tris protein gel (Bio-Rad). Protein was transferred at 4°C to nitrocellulose membrane for 2 hours at 50V. Following transfer, blocking was performed overnight at 4°C with a 5% milk solution in 1x PBS supplemented with 0.1% Tween 20. The primary antibodies used were a 1:300 dilution of antibody against the DENV2 NS3 protein (mouse polyclonal antibody raised against the DENV2 NS3 protein as described in 23), and 1:100 dilution of β-actin (rabbit polyclonal antibody, Invitrogen). The secondary antibodies used were a 1:3000 dilution of goat-anti-mouse IRDye 800CW, and goat-anti-rabbit IRDye 680RD (Li-Cor Biosciences). Blots were imaged on a ChemiDoc MP Imaging System (Bio-Rad), and quantified in ImageJ utilizing area under the curve analysis.

### Statistical analysis

In each figure and/or figure legend, statistical analysis details are noted. All results are expressed as mean values with standard deviation. Statistical significance was determined using a one-way Analysis of Variance (ANOVA) with Dunnett’s multiple comparisons test, or an unpaired t tests using Prism software version 9.0 (GraphPad Software, La Jolla, California USA).

## Acknowledgements

This work was supported by the American Society for Microbiology Undergraduate Fellowship to SAM and funding from the Department of Microbiology, Immunology and Pathology and the Office of the Vice President for Research, Colorado State University.

## Author statement

LAS, SAM, EL, NC, RCG and RP contributed to Conceptualization and development of methodology. LAS, SAM, EL, NC, RCG, GR, AN, CM and RP contributed to data acquisition and analysis. LAS, SAM, EL, NC, RCG and RP contributed to writing, reviewing and editing.

---

**Supplementary Table 1.**
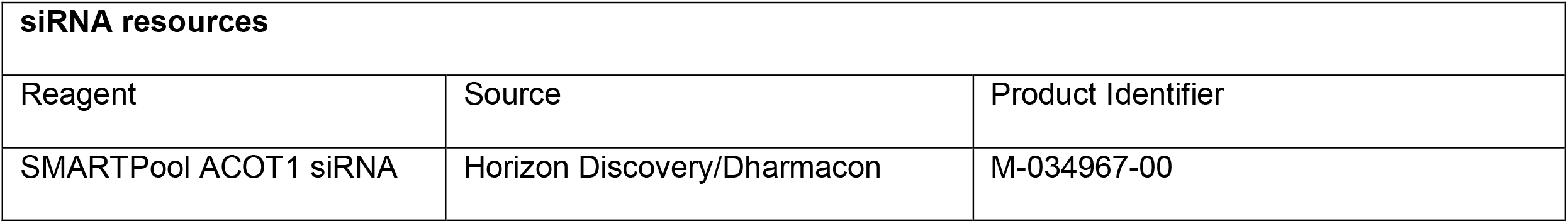
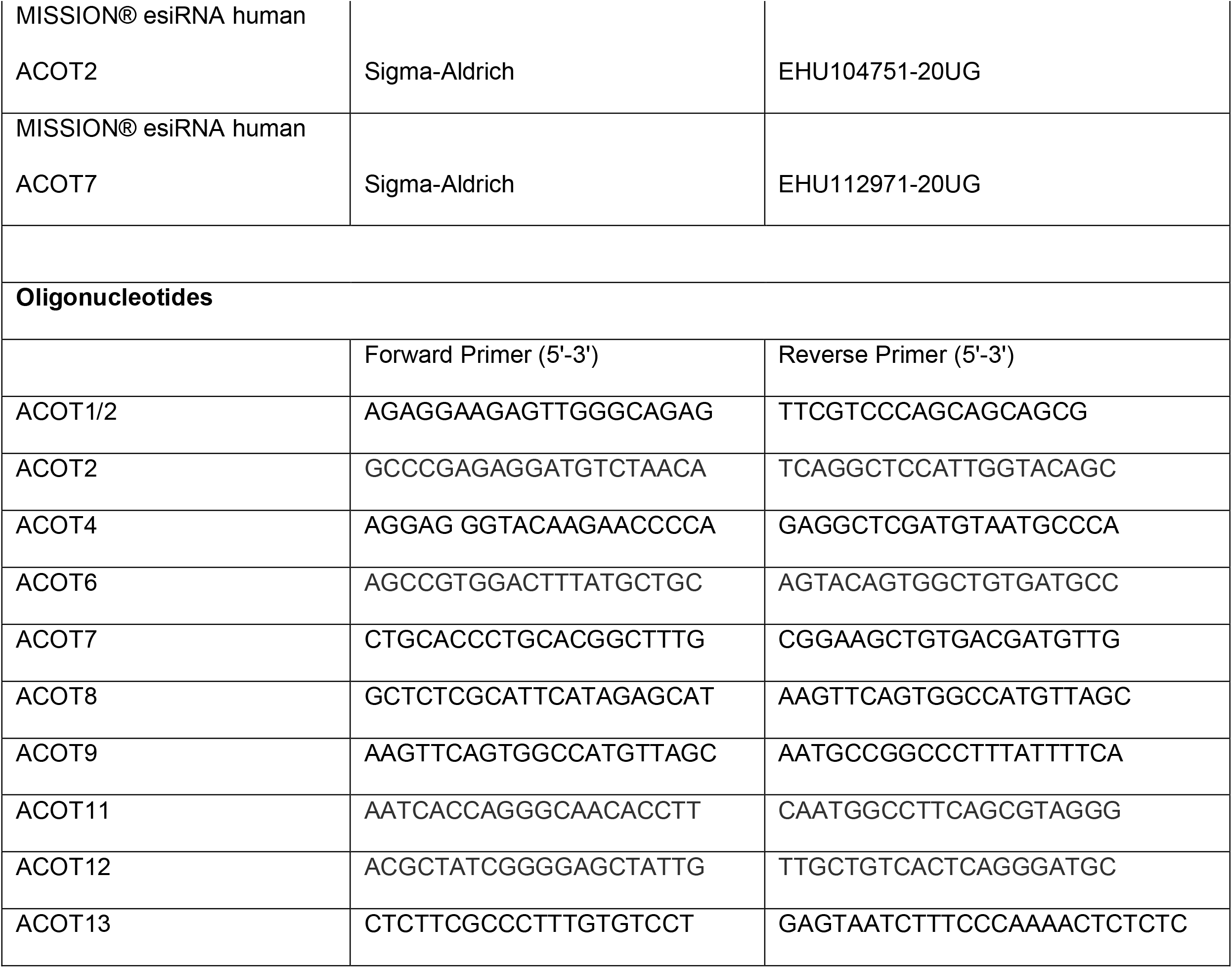
siRNA and oligonucleotide resources used in this study.

